# BIO-LGCA: a cellular automaton modelling class for analysing collective cell migration

**DOI:** 10.1101/2020.10.29.360669

**Authors:** Andreas Deutsch, Josué Manik Nava-Sedeño, Simon Syga, Haralampos Hatzikirou

## Abstract

1

Collective dynamics in multicellular systems such as biological organs and tissues plays a key role in biological development, regeneration, and pathological conditions. Collective tissue dynamics - understood as population behaviour arising from the interplay of the constituting discrete cells - can be studied with on- and off-lattice agent-based models. However, classical on-lattice agent-based models, also known as cellular automata, fail to replicate key aspects of collective migration, which is a central instance of collective behaviour in multicellular systems.

To overcome drawbacks of classical on-lattice models, we introduce an on-lattice, agent-based modelling class for collective cell migration, which we call biological lattice-gas cellular automaton (BIO-LGCA). The BIO-LGCA is characterised by synchronous time updates, and the explicit consideration of individual cell velocities. While rules in classical cellular automata are typically chosen ad hoc, rules for cell-cell and cell-environment interactions in the BIO-LGCA can also be derived from experimental cell migration data or biophysical laws for individual cell migration. We introduce elementary BIO-LGCA models of fundamental cell interactions, which may be combined in a modular fashion to model complex multicellular phenomena. We exemplify the mathematical mean-field analysis of specific BIO-LGCA models, which allows to explain collective behaviour. The first example predicts the formation of clusters in adhesively interacting cells. The second example is based on a novel BIO-LGCA combining adhesive interactions and alignment. For this model, our analysis clarifies the nature of the recently discovered invasion plasticity of breast cancer cells in heterogeneous environments. A Python package which implements various interaction rules and visualisations of BIO-LGCA model simulations we have developed is available at https://github.com/sisyga/BIO-LGCA.

**Author summary:** Pattern formation during embryonic development and pathological tissue dynamics, such as cancer invasion, emerge from individual inter-cellular interactions. In order to study the impact of single cell dynamics and cell-cell interactions on tissue behaviour, one needs to develop space-time-dependent on- or off-lattice agent-based models (ABMs), which consider the behaviour of individual cells. However, classical on-lattice agent-based models also known as cellular automata fail to replicate key aspects of collective migration, which is a central instance of collective behaviour in multicellular systems. Here, we present the rule- and lattice-based BIO-LGCA modelling class which allows for (i) rigorous derivation of rules from biophysical laws and/or experimental data, (ii) mathematical analysis of collective migration, and (iii) computationally efficient simulations.

## 3 Introduction

Systems biology and mathematical modelling is rapidly expanding its scope from the study of single cells to the analysis of collective behaviour in multicellular tissue- and organ-scale systems. In such systems, individual cells may interact with their environment (hapto- and chemotaxis, contact guidance, etc.) or with other cells (cell-cell adhesion, contact inhibiton of locomotion, etc.) and produce collective patterns exceeding the cells’ interaction range. To study collective behaviour in such systems theoretically and/or computationally, a mathematical model must be decided upon as a first step. State-continuous models describe the dynamics of cell densities. Their lack of resolution at the individual scale makes them inappropriate to investigate the role of individuals in collective behaviour. Agent-based models, on the other hand, are particularly suited to the study of collective behaviour in multicellular systems, as they resolve individual cell dynamics, and thus allow for the analysis of large-scale tissue effects of individual cell behaviour.

In the context of multicellular tissue dynamics, various agent-based models have been developed to analyse tissue dynamics as a collective phenomenon emerging from the interplay of individual biological cells. In these models, cells are regarded as separate, individual units, contrary to continuum models, which neglect the discrete individual cell nature, and where tissue dynamics is derived from conservation and constitutive laws, drawing parallels to physical systems. Since agent-based models represent individual biological cells, distinct cell phenotypes can be taken into account, which may be fundamental for analysing the organisation at the tissue level. For example, it has been shown that cell-to-cell variability plays a key role in tumour progression and resistance to treatment [1]. Moreover, with the advance of high performance computing, agent-based models can be used to analyse *in vitro* systems at a 1:1 basis even for large cell population sizes.

Agent-based models can be classified into on-lattice and off-lattice or “lattice-free” models depending on whether or not cell movement is restricted to an underlying lattice (see [2] for references). Various off-lattice models exist to study different types of single and collective cell migration [3, 4, 5, 6, 7, 8].

In lattice models, either (i) a lattice site may be occupied by many biological cells (e.g., [9]), (ii) a site may be occupied by at most one single biological cell, or (iii) several neighbouring lattice sites may represent a single biological cell (e.g., [10]). Model types (i) and (ii) can mimic volume exclusion effects, (iii) can qualitatively capture cell deformation and compression, while each of the three approaches can describe the effects of mechanical forces of one cell on its neighbour, or on a group of neighbouring cells to some extent.

Lattice models are equivalent to cellular automata (CA), which were introduced by J. v. Neumann and S. Ulam in the 1950s as models for individual self-reproduction [11]. A cellular automaton consists of a regular spatial lattice in which each lattice node can assume a discrete, typically finite number of states. The next state of a node solely depends on the states in neighbouring sites and a deterministic or stochastic transition function. One distinguishes CA with synchronous and asynchronous update. Cellular automata provide simple models of self-organising systems in which collective behaviour emerges from an ensemble of interacting “simple” components - being it molecules, cells or organisms [12, 13, 14]. The interacting particle system (IPS) is an example of probabilistic CA with asynchronous update. Here, proliferation, death, and migration of biological cells are modeled as stochastic processes.

However, when modelling collective migration phenomena, classical CA and, particularly, IPS models have major drawbacks which are due to the strict volume exclusion and asynchronous update. Most importantly, these models fail to reproduce collective movement at unit density, since volume exclusion at high densities results in a “jammed state”. However, a fluidised state at unit density is an important case of collective migration especially in epithelial tissues. Furthermore, the asynchronous update in IPS models may lead to oscillating density spikes. For example, in an IPS model for persistent motion in a crowded environment, cells at the invasion front detach and leave gaps behind that are subsequently filled by following cells [15]. This is an artefact of the asynchronous update since invasion can happen while cells stay connected. Moreover, classic CA models consider only cell position and not explicitly cell momentum, complicating the modelling of collective cell migration mediated primarily through changes in momentum, rather than density [16].

The lattice-gas cellular automaton (BIO-LGCA) introduced here, is a cellular automaton in which lattice sites are updated synchronously, and which explicitly considers individual cell velocities. These features make the BIO-LGCA appropriate for modelling collective migration phenomena where cell interactions result in directional changes of velocity, and where high cell densities do not hamper movement.

The structure of the paper is as follows: we first formally define the BIO-LGCA model class. Then, we construct *biophysical BIO-LGCA rules* from microscopic Langevin models for selected cases of single and collective cell migration. Subsequently, we demonstrate how to generate *data-driven BIO-LGCA rules* from experimental single cell migration data (Fig. 1). Furthermore, we show that, in specific cases, the biophysical and the data-driven approaches converge to the same functional form. For this case, we introduce several biologically relevant model examples. Finally, we present two examples of mean-field analysis. The first example allows to predict the formation of cluster patterns in simulations of a BIO-LGCA model with adhesive cell-cell interaction. The second example analyses a novel BIO-LGCA combining adhesive interactions and alignment to explain the recently discovered invasion plasticity of breast cancer cells in heterogeneous environments. We end with a critical discussion of the BIO-LGCA modelling framework.

**Figure 1:**
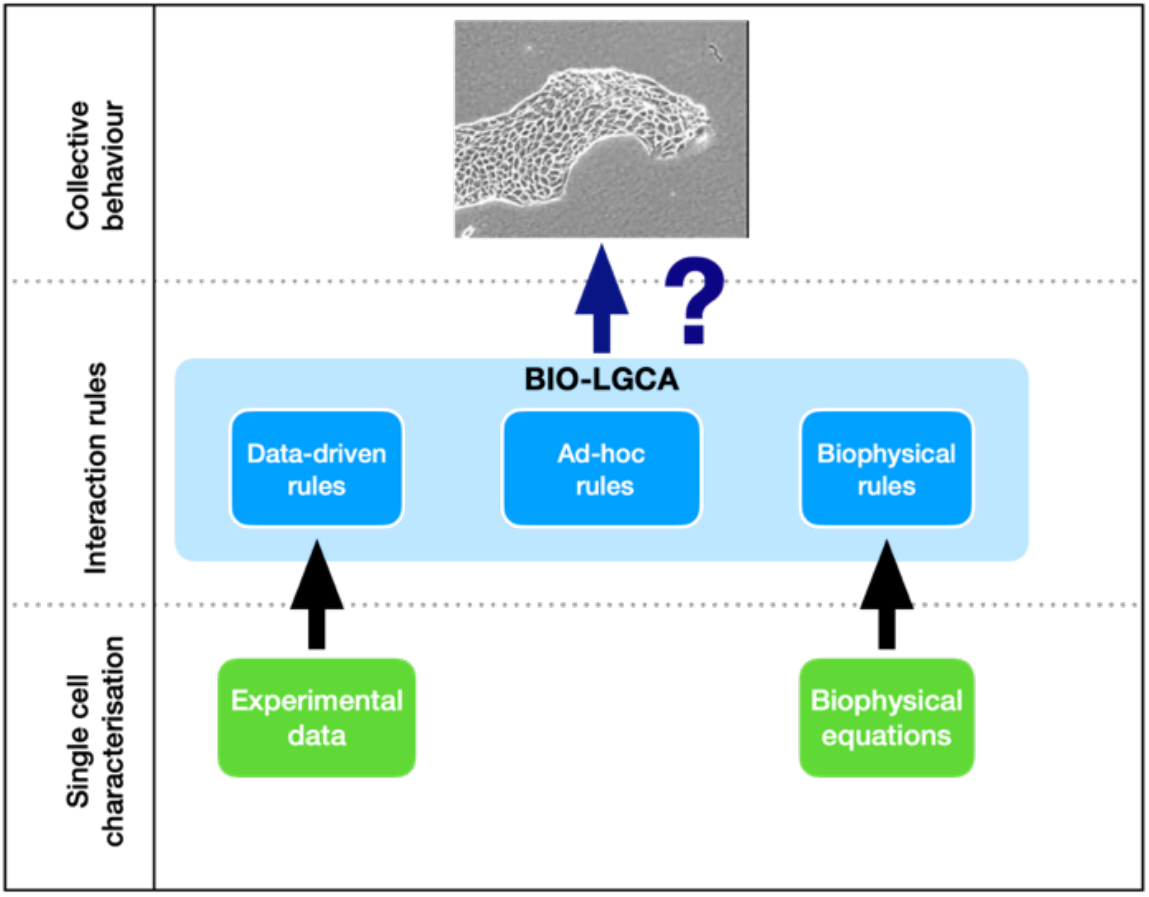
BIO-LGCA modelling: The key question is to extract the interaction rules underlying a particular collective phenomenon in a population of cells. BIO-LGCA interaction rules can be chosen ad hoc, extracted from experimental single cell migration data, or derived from biophysical equations for single cell migration.

## 4 Materials and Methods

A BIO-LGCA is defined by a discrete spatial lattice *ℒ*, a discrete state space ϵ, a neighbourhood 𝒩 and local rule-based dynamics.

### 4.1 Lattice

The regular lattice *ℒ* ⊂ ℝ^*d*^ consists of nodes **r** ∈ *ℒ*. Every node has *b* nearest-neighbours, where *b* depends on the lattice geometry. Each lattice node **r** ∈ *ℒ* is connected to its nearest neighbours by unit vectors **c**_*i*_, *i* = 1, …, *b*, called velocity channels. In addition, a variable number *a* ∈ ℕ_0_ of rest channels (zero-velocity channels) **c**_*j*_ = **0**, *b* < *j* ≤ *a* + *b*, is allowed (Fig. 2). The parameter *K* = *a* + *b* defines the maximum *node capacity*.

**Figure 2:**
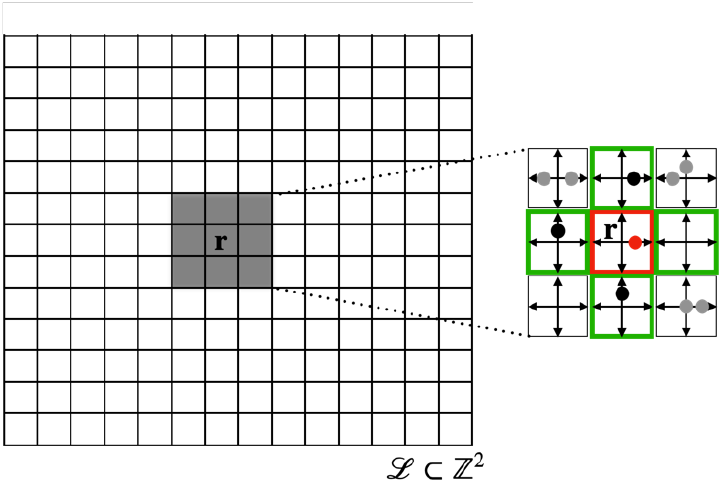
Lattice and neighbourhood in the BIO-LGCA: example of square lattice (left). The node state is represented by the occupation of velocity channels (right); in the example, there are four velocity channels **c**_1_, **c**_2_, **c**_3_, **c**_4_, corresponding to the lattice directions, and one “rest channel” **c**_0_. Filled dots denote the presence of a cell in the respective velocity channel; right: von Neumann neighbourhood (green) of the red node.

### 4.2 Neighbourhood

The set 𝒩, the neighbourhood template, defines the nodes which determine the dynamics of the node 0 ∈ *ℒ*. Throughout this work, the neighbourhood will be assumed to be a von Neumann neighbourhood (Fig. 2), defined as 𝒩^*b*^ := 𝒩^*b*^(0) = {**c**_**1**_, **c**_**2**_, …, **c**_**b**_}, but other neighbourhood choices are possible. In general, 𝒩 (**r**) := 𝒩^*b*^(**r**) = 𝒩^*b*^ + **r**, specifies the set of lattice nodes which determine the dynamics of the state at node **r** ∈ *ℒ*. ^1^

### 4.3 State space

The state space in LGCA is defined through the occupation numbers *s*_*j*_ ∈ {0, 1}, *j* = 1, …, *K*. These occupation numbers represent the presence (*s*_*j*_ = 1) or absence (*s*_*j*_ = 0) of a cell in the channel **c**_*j*_ within some node. Then, the configuration of a node is given by the state vector

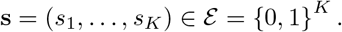

This reflects an *exclusion principle* which allows not more than one cell at the same node within the same channel simultaneously. As a consequence, each node **r** ∈ *ℒ* can host up to *K* cells, which are distributed in different channels.

It is possible to consider more than one cell phenotype in the BIO-LGCA model. In this case each phenotype is indexed by *σ* ∈ Σ ⊂ N. Then, the configuration vector is given by

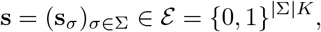

where | · | denotes the cardinality of a set. Each node will be able to support up to |Σ|*K* cells.

Two useful quantities for a given node are the *total number of cells at the node n*(**s**) and the *momentum/node flux* **J**(**s**), defined as

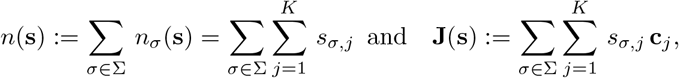

where *n*_*σ*_(**s**) is the *σ* number of cell phenotypes.

### 4.4 Dynamics

In general, in cellular automata a new lattice configuration is created according to a local rule that determines the new state of each node in terms of the current states of the node and the nodes in its neighbourhood. In order to determine a new lattice configuration, the local rule is applied independently and simultaneously at every node **r** of the lattice. Mathematically, in probabilistic cellular automata, the local rule can be interpreted as a transition probability *P* (**s** → **s**′) to replace a current configuration **s** with a new node configuration **s**′.

In a BIO-LGCA, local rules are composed of a particular combination of operators for stochastic reorientation (*𝒪*), phenotypic switching (*𝒮*), and stochastic cell birth and death (*ℛ*), as well as a deterministic propagation operator (*𝒫*) (see figure 3). The propagation and reorientation operators together define cell movement, while phenotypic switching allows cells to stochastically and reversibly transition between phenotypes. In a BIO-LGCA, the stochastic operators are applied sequentially to every node, such that the transition probability can be expressed as

**Figure 3:**
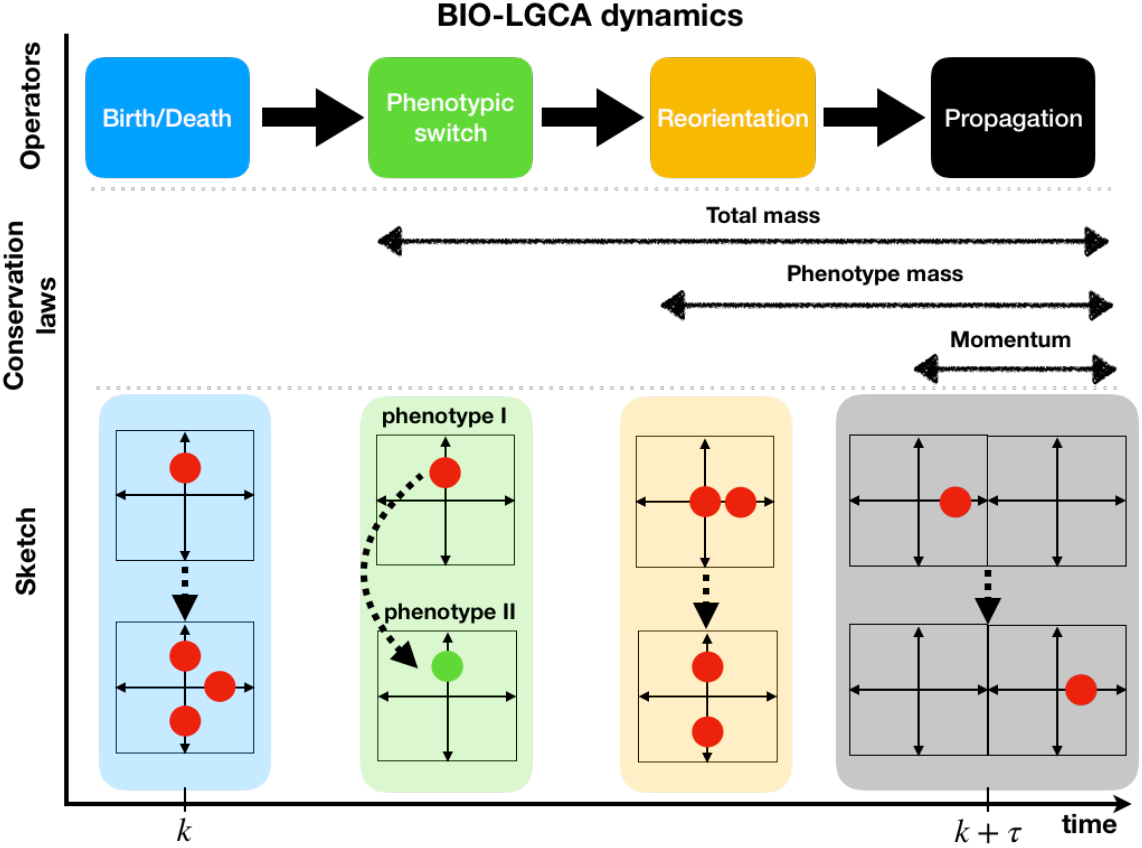
Operator-based dynamics of the BIO-LGCA: Propagation *𝒫*, reorientation *𝒪*, phenotypic switch *𝒮*, and birth/death operators *ℛ* (top); conservation laws maintained by the different operators (middle); sketches of the operator dynamics (bottom), see text for explanations.

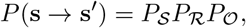

where *P*_*i*_, *i* ∈ {*𝒮, ℛ, 𝒪*}are the transition probabilities of the corresponding operator. In this way, a post-interaction node configuration **s**′ is defined as the resulting node configuration after subsequent application of the stochastic operators, i.e. **s**′ = **s**^*𝒫*∘ *ℛ*∘ *𝒪*^. Subsequently, the deterministic propagation operator *𝒫* is applied: cells occupying velocity channels at the node, i.e. moving cells, are translocated to neighbouring nodes in the direction of their respective velocity channels. The time step increases once the propagator operator has been applied. Accordingly, the dynamics of the BIO-LGCA can be summarized in the stochastic microdynamical equation

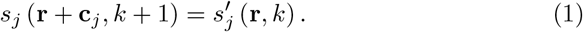

### 4.5 Simulator

https://imc.zih.tu-dresden.de//biolgca/ https://github.com/sisyga/BIO-LGCA

## 5 Results

### 5.1 BIO-LGCA rule derivation

In classical cellular automata, transition probabilities are typically chosen *ad hoc*. Here, we show that BIO-LGCA rules can also be derived from biophysical equations of motion and from experimental data. In the following, we disregard birth/death processes and phenotypic transitions. Accordingly, the corresponding BIO-LGCA model is specified exclusively by its reorientation probability (cp. 4.4 and fig. 3). We first present a method to derive the BIO-LGCA reorientation probability from a Langevin model for cell migration [17]. A similar procedure has also been used to derive rules for CA models describing molecular movement in crowded environments [18]. Secondly, we sketch a method to obtain the reorientation probability from experimental observations [19]. In certain cases, independent of the applied rule derivation method, the functional form of the transition probability will be the same (Fig. 4).

**Figure 4:**
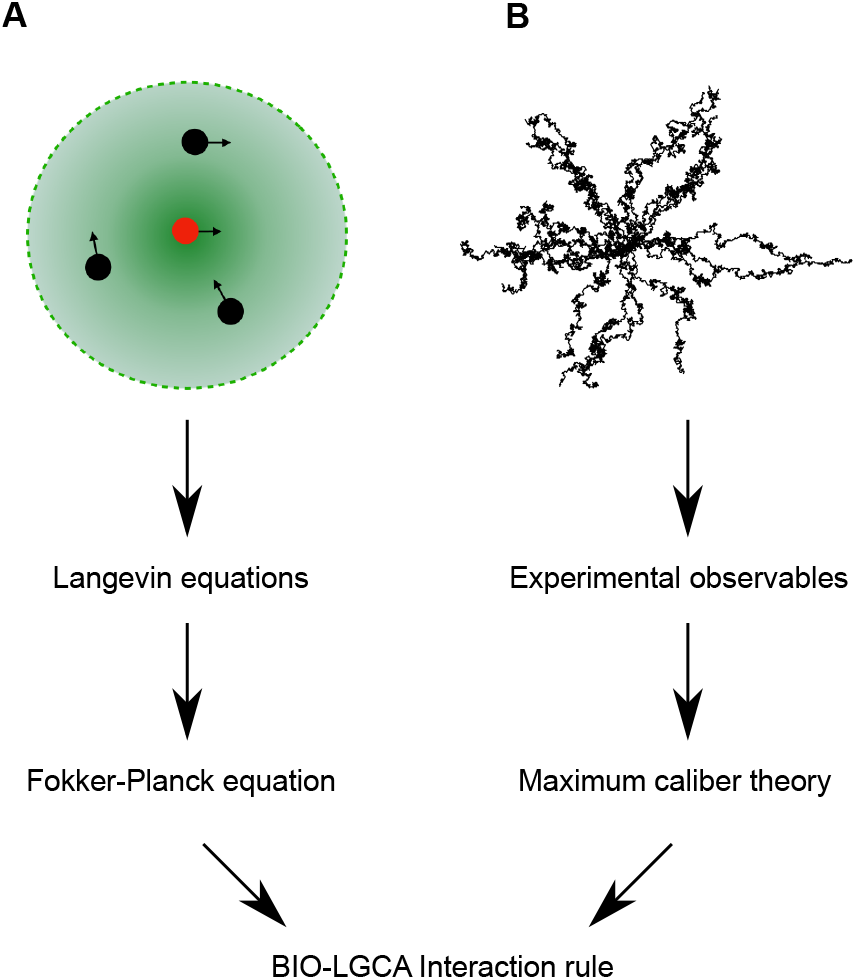
Rule generation in BIO-LGCA models. (A) Starting from the Langevin equations of a self-propelled particle model, the interaction rule can be obtained from the steady-state distribution of the associated Fokker-Planck equation. (B) Alternatively, experimental observables can be used as input for the maximum caliber theory to derive the probabilities of cell tracks and the corresponding interaction rule. Image credit: [23].

**Figure 5:**
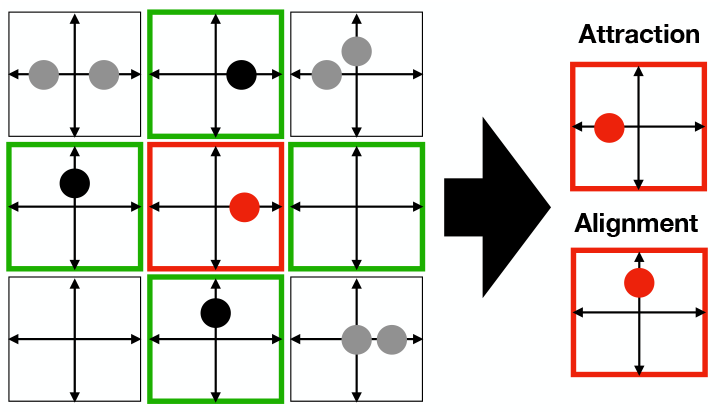
Basic interactions with neighbourhood impact; node configuration (red) before and after application of stochastic interaction rule: cell-cell attraction (top), cell alignment (bottom).

**Figure 6:**
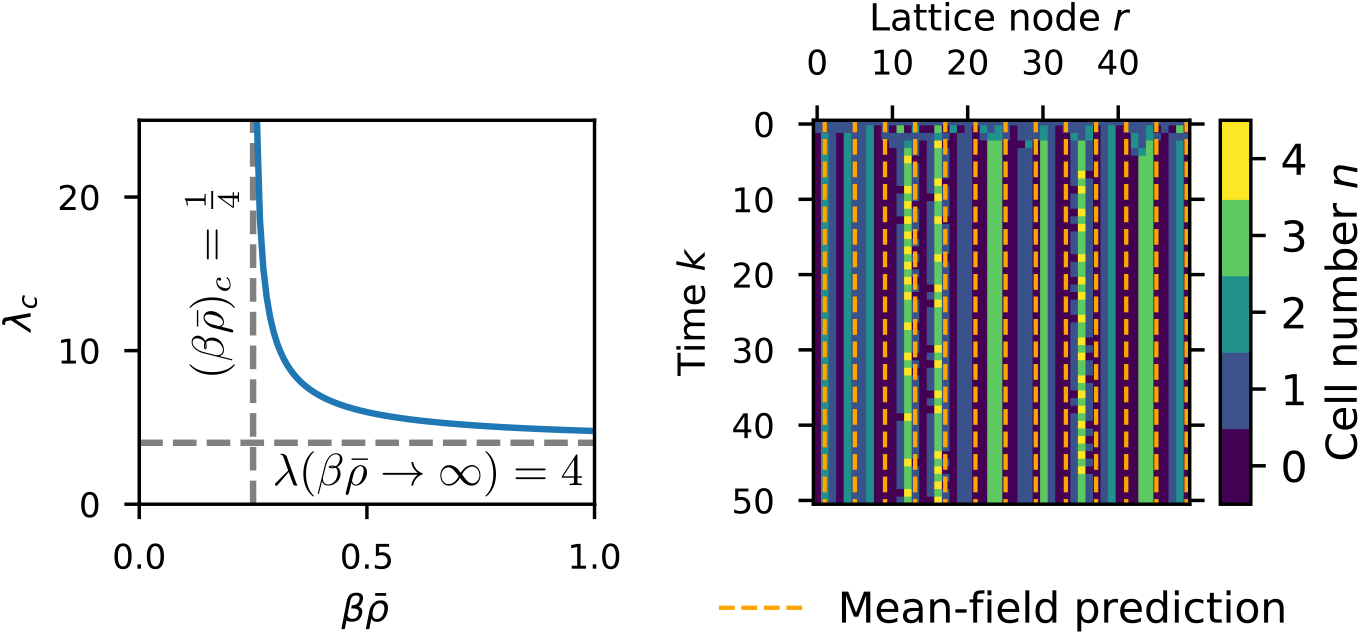
Pattern formation in the LGCA aggregation model. Left: critical wave length obtained from the mean-field analysis. The critical wave length diverges for 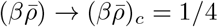, and 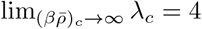. Right: emergence of a periodic pattern from a homogeneous initial state. The horizontal distance of the orange dashed lines is equal to the critical wave length predicted by mean-field analysis. Parameters: *β* = 100, 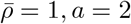.

#### 5.1.1 Reorientation dynamics derived from biophysical equations of motion

It has been shown that various types of cell migration can be described by self-propelled particle models (SPPs). These off-lattice models are defined by a set of stochastic differential equations governing the motion of discrete cells in an overdamped situation, e.g in a highly viscous medium. The stochastic differential equations encoding individual cell motion (as introduced in [20]) are Langevin equations, where a stochastic variable *θ*_*m*_ describes the orientation of the *m*-th cell in the system which moves with a constant speed *v*_0_ ∈ ℝ+and orientation *θ*_*m*_(*t*) ∈ [0, 2*π*) varying according to a given interaction potential and influenced by noise. The Langevin equations of motion read [21]

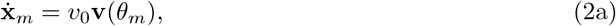

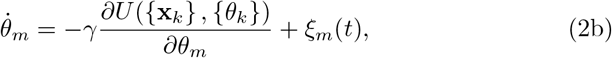

where **x**_*m*_ ∈ ℝ^*d*^is the cell’s spatial position, **v**(*θ*_*m*_) ∈ ℝ^*d*^ is a unit vector pointing in the direction of the cell’s displacement, *γ* ∈ ℝ+is a relaxation constant, and *ξ*_*m*_(*t*) is a white noise term with zero mean and correlation ⟨ *ξ*_*m*_ (*t*_1_) *ξ*_*n*_ (*t*_2_) ⟩ = 2*D*_*θ*_*δ* (*t*_2_ − *t*_1_) *δ*_*m,n*_. The heart of the model is the potential *U* ({**x**_*k*_}, {*θ*_*k*_}): ℝ^*Nd*^ × [0, 2*π*)^*N*^ ↦ ℝ, where *N* is the number of cells within the *m* th cell’s neighbourhood of interaction, and {**x**_*k*_} and {*θ*_*k*_}are the sets of all neighboring cells’ positions and orientations, respectively. This potential encodes the biophysical mechanisms that dictate the cell’s reorientation. Here, we assume that the reorientation potential only depends on the orientations of neighbouring cells, though a dependence on cell positions is also possible [22].

The probability density function of the stochastic variable *θ*_*m*_ governed by the Langevin equations is given by the Fokker-Planck equation

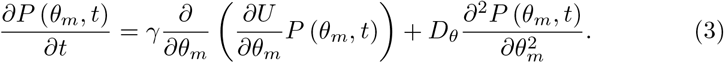

If we assume fast relaxation times for the solution of the Fokker-Planck equation, then one can choose the stationary solution as the probability of cell *m* to have an orientation *θ*_*m*_, i.e.

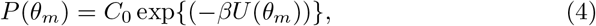

where *β* = *γ/D*_*θ*_ and *C*_0_ is an integration constant. Applying a discretisation of the particle orientations and under the assumption that the relaxation time of the Fokker-Planck solution is smaller than the BIO-LGCA time step and that the channel occupations in the LGCA model are independent one can derive the reorientation probability

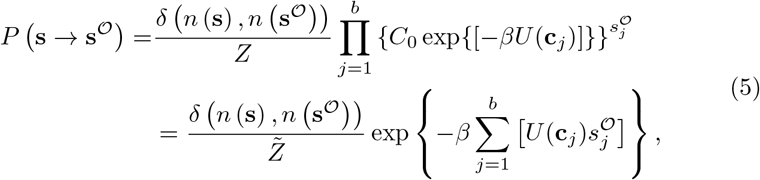

where 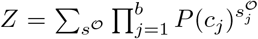 and 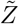 is a normalisation constant in which the integration constant *C*_0_ has been absorbed [17].

##### Collective migration

SPP models for collective cell migration often use interaction potentials of the following form

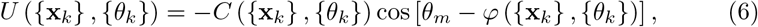

where amplitude *C* (interaction strength) and shift *ϕ* (optimal orientation) may depend on the positions and/or orientations of all cells within the neighbourhood of interaction (including the central cell). Using trigonometric identities, the reorientation potential can then be rewritten as

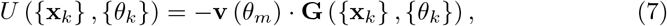

where **G** ({**x**_*k*_}, {*θ*_*k*_}) is called the *local director field*, whose norm and argument are, respectively, ∥**G** ({**x**_*k*_}, {*θ*_*k*_})∥ = *C* ({**x**_*k*_}, {*θ*_*k*_}) and arg [**G** ({**x**_*k*_}, {*θ*_*k*_})] = *ϕ* ({**x**_*k*_}, {*θ*_*k*_}). Substituting Eq. 7 in Eq. 5, and using the linearity of the internal product, the transition probability of the reorientation operator is

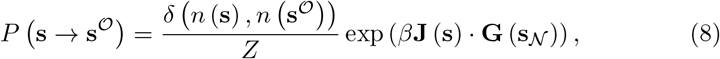

where **G** (**s**_𝒩_) is the local director field of the neighbourhood configuration, and **J** (**s**) is the node flux, as described previously.

More generally, whenever the reorientation potential can be expressed as

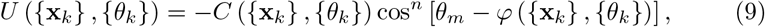

with *n* ∈ ℕ, the argument of the exponential in the transition probability can be expressed as an internal product of two vectors. In the specific case of *n* = 2, using trigonometric functions, one can arrive at the transition probability

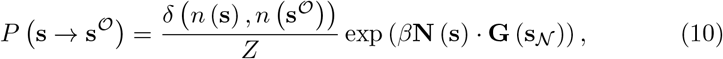

where **N** (**s**) is the *local nematic alignment vector*, and is defined as

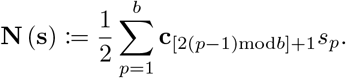

Thus, the reorientation probabilities has the same general form whenever the interaction potential is conservative (i.e. time-independent) and consists of a pairwise comparison between the angles and/or positions of neighbouring cells.

#### 5.1.2 Data-driven rules

Besides from Langevin equations defining SPP models, it is also possible to derive BIO-LGCA reorientation probabilities from experimental data (Fig. 4). For this, we assume that certain observables, e.g. an autocorrelation function, have been obtained from primary migration data. Then, we can use the maximum caliber (or maximum path entropy) formalism [24] to obtain the most unbiased probability distribution of paths that reproduces the experimental observables. This translates into maximising the following functional

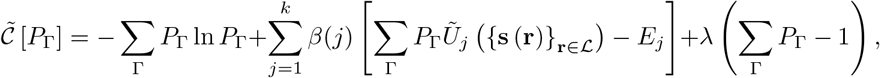

where *P*_Γ_ is the probability of a cell to follow a certain spatial trajectory Γ, *β*(*j*) and *λ* are Lagrange multipliers, *Ũ* _*j*_ {**s** (**r**)}_**r**∈*ℒ*_ is the value of the observable at time step *j* depending on the configuration of the lattice {**s** (**r**)} _**r**∈*ℒ*_, and *E*_*j*_ is the value of the experimental observable at time step *j*. The first term of the functional is the entropy, which we want to maximise. The second term restricts the resulting probabilities to match the experimental observation. Since we assume the experimental observable, *E*_*j*_ to be a time-dependent function, a Lagrange multiplier *β*(*j*) is needed for every time step *j*. The last term guarantees the normalisation of probabilities, which requires an additional Lagrange multiplier, *λ*.

The optimisation of this functional yields an optimal value for the path probabilities 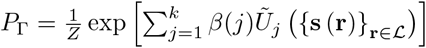, where the value of *β*(*j*) is such that *E*_*k*_ =Σ Γ *P*_Γ_*Ũ* _*j*_ ({**s** (**r**)} _**r**∈*ℒ*_).

If the process is Markovian, then one may decompose the path probability into individual channel occupation probabilities for each time step *k*, as

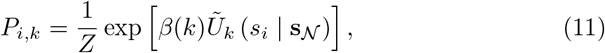

where *Z* is a normalisation constant and the observable is dependent on the occupancy of the *i*-th channel of the node and conditioned to a certain configuration of its interaction neighbourhood. For example, if the observation is the autocorrelation function *g*(*t*) = ⟨ **v**_0_ · **v**_*t*_ ⟩, where **v**_*t*_ denotes the normalized velocity of a cell at time *t*, determined from experimental data, then the corresponding channel occupation probabilities are found to be

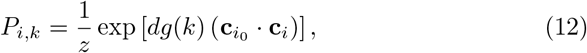

where *z* is the normalisation constant for the transition probability, *d* is the dimension of space, and 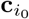 is the initial orientation of the cell.

If we assume independence among cells within the same node, we arrive at a reorientation probability of the form

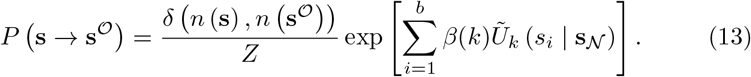

Note that, if both observation and observable are time-independent, then the Lagrange multiplier *β*(*k*) = *β* is also time independent, and the transition probabilities are given by

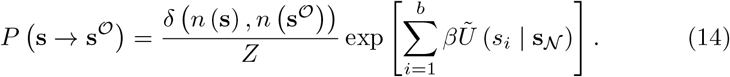

Furthermore, if *Ũ* (*s*_*i*_ |**s**_𝒩_) = *s*_*i*_**c**_*i*_ · **G** (**s**_𝒩_), then Eq. 14 reduces to Eq. 8. In conclusion, we can construct rules of the BIO-LGCA directly from experimental observables and the structure of these rules is the same as in our ad-hoc and SPP model-derived rules.

### 5.2 BIO-LGCA rules for single and collective cell migration

Here, we present key examples of transition probabilities corresponding to reorientation operators, which model important elementary single-cell and collective behaviours. Note that several of these examples’ transition probabilities have the general form of Eq. 8.

#### 5.2.1 Single cell migration

##### Random walk

Random walks are performed by cells such as bacteria and amoebae in the absence of any environmental cues. Random walk of cells can be modeled by a reorientation operator with the following transition probabilities:

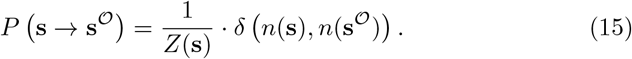

This rule conserves mass, i.e. cell number.

##### Chemotaxis

Chemotaxis describes the dependence of individual cell movement on a chemical signal gradient field. Accordingly, spatio-temporal pattern formation at the level of cells and chemical signals can be observed. Chemotactic patterns result from the coupling of different spatio-temporal scales at the cell and the molecular level, respectively.

To mimick a chemotactic response to the local signal concentration, we define the signal gradient field

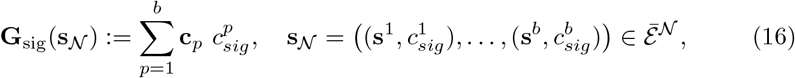

Where 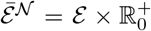. Chemotaxis can be modeled through a reorientation operator with transition probabilities given by

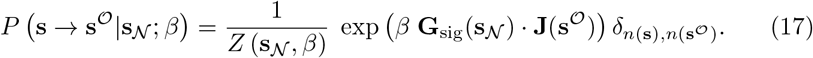

where *β* is the *chemotactic sensitivity* of the cells.

With large probability, cells will move in the direction of the external chemical gradient **G**_sig_.

##### Haptotaxis

We consider cell migration in a static environment that conveys directional information expressed by a vector field

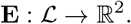

A biologically relevant example is haptotactic cell motion of cells responding to fixed local concentration differences of adhesion molecules along the extracellular matrix (ECM). In this example, the local spatial concentration differences of integrin ligands in the ECM constitute a gradient field that creates a “drift” **E** [25].

The transition probabilities associated to the reorientation operator, given a vector **E** ∈ ℝ^2^, is given by

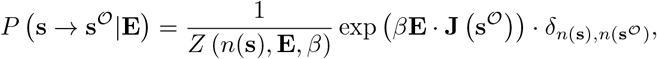

Where **E** ∈ ℝ^2^

In this case, cells preferably move in the direction of the external gradient **E**.

##### Contact guidance

We now focus on cell migration in environments that convey orientational, rather than directional, guidance. Examples of such motion are provided by neutrophil or leukocyte movement through the pores of the ECM, the motion of cells along fibrillar tissues, or the motion of glioma cells along fiber track structures. Such an environment can be represented by a second rank tensor field that encodes the spatial anisotropy along the tissue. In each point, the corresponding tensor informs the cells about the local orientation and strength of the anisotropy and induces a principal (local) axis of movement. Thus, the enviroment can again be represented by a vector field

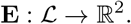

Contact guidance can be modeled through a reorientation operator with transition probabilities defined as

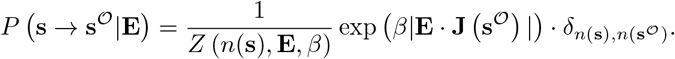

#### 5.2.2 Collective cell migration

Several kinds of organisms, as well as biological cells, e.g. fibroblasts, can align their velocities globally through local interactions. Here, we introduce a reorientation operator where the local director field is a function of the states of several channels and nodes, reflecting the influence of neighbouring cells during collective cell migration.

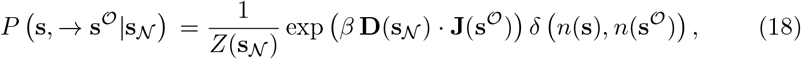

where 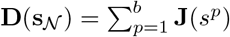 is the *local cell momentum* This particular reorientation probability triggers cell alignment [26].

#### 5.2.3 Attractive interaction

Biological cells can interact via cell-cell adhesion, through filpodia cadherin interaction, for example. Agent attraction/adhesion can be modeled with a reorientation operator with the following probability distribution.

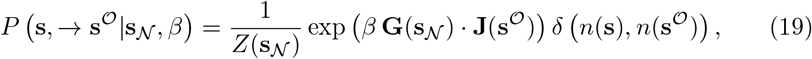

where 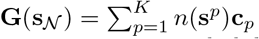 is the *density gradient field*.

This reorientation probability favors cell agglomeration. A similar rule has been introduced in [27].

### 5.3 Mean-field analysis of a BIO-LGCA model with attractive interaction

We here demonstrate the mean-field analysis of the BIO-LGCA model attractive interaction (cp. eq. (19)). This analysis allows to predict collective behaviour in the form of cell aggregation. In particular, we calculate the critical sensitivity *β*_*c*_, such that aggregation occurs for *β > β*_*c*_, while a homogeneous initial conditions is stable for *β* < *β*_*c*_. Under “mean-field” we understand that we neglect correlations between the occupation numbers of different channels and that we approximate the mean value of any function *f* of a random variable *X* by the function evaluated at the mean value of the random variable, i.e. ⟨ *f* (*X*) ⟩ ≈ *f* (⟨ *X*⟩). As we are interested in the onset of aggregation from a homogeneous initial state with low density 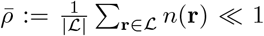 and weak interaction *β* ≪ 1 we can linearise the transition probabilities. We further assume that there is at most one cell at each node and therefore only consider single-cell transitions (*n* = 1). For the partition function *Z* we then obtain

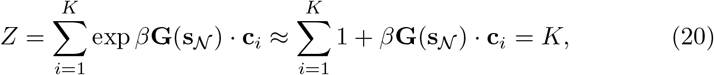

due to the lattice symmetry. For the single-cell transition probability we obtain

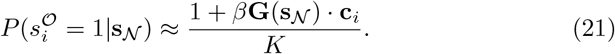

Since the transition probability only depends on the number of cells on the neighbouring nodes, but not on their distribution on the channels, we analyse the mean local density

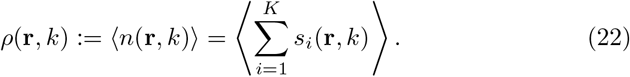

According to the propagation rule the cell number *n*(*r, k* + 1) is given by

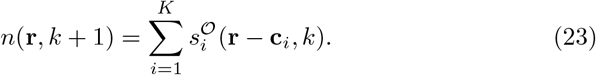

We calculate the expected value under the mean-field assumption in terms of numbers of cells *n*(**r**) at node **r** ∈ *ℒ*, and the number of cells **n**_N(**r**)_ in the neighbourhood of **r** ∈ *ℒ* as

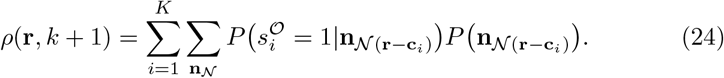

As in the low-density regime *P* (*n*(**r**) *>* 1) ≈0 ∀**r** ∈ *ℒ*, we can use the single-cell transition probability eq. (21) and the factorizing probability distribution under our mean-field assumption to obtain

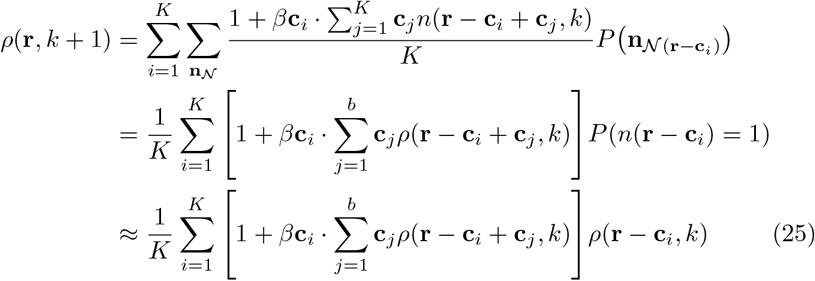

To proceed, we assume a one-dimensional lattice, where *b* = 2, *c*_1,2_ = ±1 with *a* rest channels to obtain the finite-difference equation (FDE)

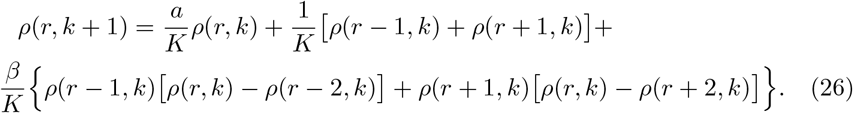

This FDE can be analysed by means of linear stability analysis. To do this, we first rewrite eq. (26) in terms of the density difference Δ*ρ*(*r, k*) := *ρ*(*r, k* + 1) − *ρ*(*r, k*), which we linearise around the steady state for a small perturbation of the form 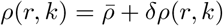

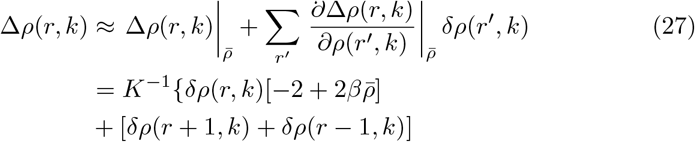

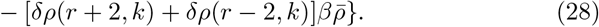

We now apply the discrete Fourier transform

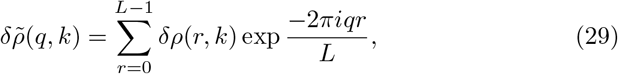

and obtain the mode-dependent FDE

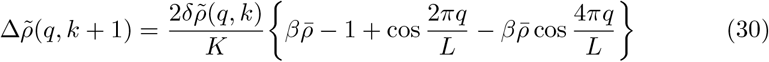

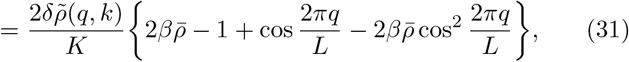

using 2 cos *x* = *e*^*ix*^ + *e*^−*ix*^ and cos 2*x* = 2 cos^2^ *x* − 1. Note that the system becomes unstable when the r.h.s. of the equation is larger than 0, meaning the perturbation grows, while it is stable with a decreasing perturbation if it is smaller than 0. To find the dominant Fourier mode *q* that maximizes the r.h.s. we assume an infinite lattice *L* → ∞ such that we can define the quasi-continuous wave number 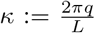 and use the derivative with respect to *κ* to calculate the maxima of the bracket on the r.h.s. of eq. (31),

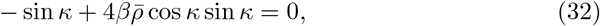

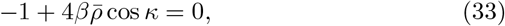

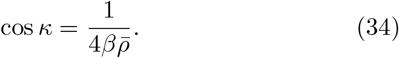

Note that we divided by sin *κ* here, neglecting the trivial solutions *κ* = 0, *π*. Clearly the solution cos 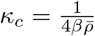 is only valid for 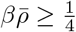 and it is the dominant wave number in this case. This in turn allows us to define the critical parameter combination 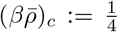. We can also calculate the dominant wave length in dependence of 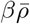 as 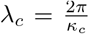, which diverges at 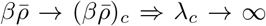 and approaches *λ*_*c*_ →4 for 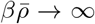. In conclusion, our mean-field analysis allows us to predict the onset of the instability of the homogeneous steady state in dependence on density *ρ* and sensitivity *β* as well as the wave length of the observed patterns without the need for computer simulations.

### 5.4 Explaining invasion plasticity of breast cancer

Progression of cancer depends on changes at the individual cell level and interactions of healthy and malignant cells. Various CA models have been suggested for selected aspects of cancer progression (see e.g. [28, 29]). Here, we shed light on the invasion plasticity of growing tumours. Solid tumours have been observed to switch between jammed (immobile, glass-like), highly correlated collective movement (active nematic phase) and single-cell-disseminating, uncorrelated (gas-like) states, which is often referred to as invasion plasticity [30, 31, 32]. These behaviours result from an interplay between cell-cell and cell-ECM interactions, which have been recently studied using a BIO-LGCA model [16]. Computational simulations and calculation of several observables showed remarkable similarity to the experimental mammary gland carcinomas. Here, we generalise the computational results in [16] by deriving an analytical theory for the observed phase transitions.

We will consider a 1D system (akin to cells moving inside an ECM duct). In this case, cells can only move in the positive or negative direction, thus the velocity channels are given by *c*_1_ = 1 and *c*_2_ = −1. As in the original model, we incorporate the effect of cell-cell adhesion as attractive and velocity alignment interactions. Additionally, we mimic steric interactions between cells by an increased tendency of cells to remain in rest channels if their neighbours occupy rest channels as well. However, while before the relative strenghts of these interacitons were assumed to be fixed, and their absolute strengths were varied by a single parameter, we here treat these mechanisms as independent, to obtain the full spectrum of possible migration modes. Our model considers *b* = 2 velocity channels and *a* rest channels, for a total of *K* = 2 + *a* channels. Under these assumptions, the BIO-LGCA transition probabilities are given by

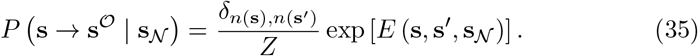

The energy function is divided into three contributions

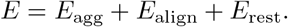

The aggregation energy is given by the logistic density gradient, *E*_agg_ = *β*_agg_*j*(**s**^*𝒪*^) *g*_agg_ (**s**_𝒩_), where 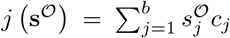 is the post-interaction flux, and the logistic density gradient

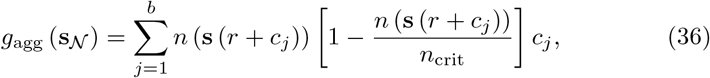

where *n*_crit_ is a parameter controlling the maximum density of cell aggregates, thus modelling the homeostatic cell density.

The alignment energy corresponds to the collective cell migration interaction (Eq. 18), here given by *E*_align_ = *β*_align_ *j* (**s**^*𝒪*^) *g*_align_ (**s**_𝒩_), where the post-interaction flux is as before and the neighbourhood flux

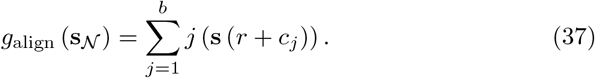

Finally, the resting energy is given by *E*_rest_ = *β*_rest_*n*_rest_ (**s**^*𝒪*^) *n*_rest_ (**s**_𝒩_), where the local resting cell density is 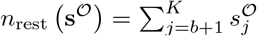 and the neighbourhood resting cell density is 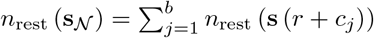.

To facilitate further analysis, we assume that the density is low, and that there is at most one cell at each node. Then, under the mean-field approximation the dynamics of the BIO-LGCA model is given by the lattice-Boltzmann (LBE) equation

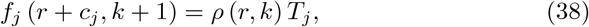

where 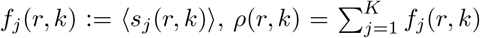, and 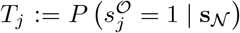 is the single particle probability given by

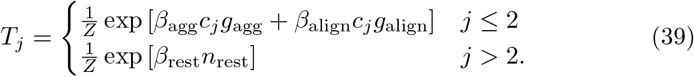

The homogeneous steady state of the LBE, where the density, 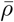, occupation of velocity channels, *f*_*v*_, and rest channels, *f*_*r*_, is constant and identical in every lattice node is given by

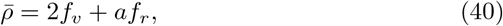

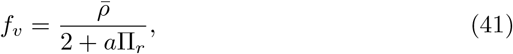

where

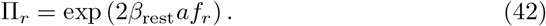

The steady states can now be calculated numerically. We linearise the LBE around the numerical steady states, and apply a discrete Fourier transform with respect to the spatial coordinates to eliminate dependencies on spatial increments. Then, the stability of the steady states is given by the eigenvalues, *λ*, of the Boltzmann propagator matrix

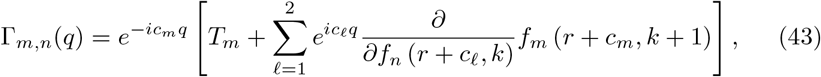

where *q* ∈ [0, 2*π*) is the wavevector, and *f*_*m*_ (*r* + *c*_*m*_, *k* + 1) is given by the LBE.

The maximum of the modulus of the Boltzmann propagator eigenvalues, |*λ*| gives information on the wavelength 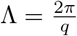 of spatial patterns observed in the model, while the argument of the eigenvalues, arg(*λ*), defines the propagation velocity of these patterns. Calculating these numerically, we find the phase space shown in Fig. 7. We can identify four distinct regions in the parameter space.

**Figure 7:**
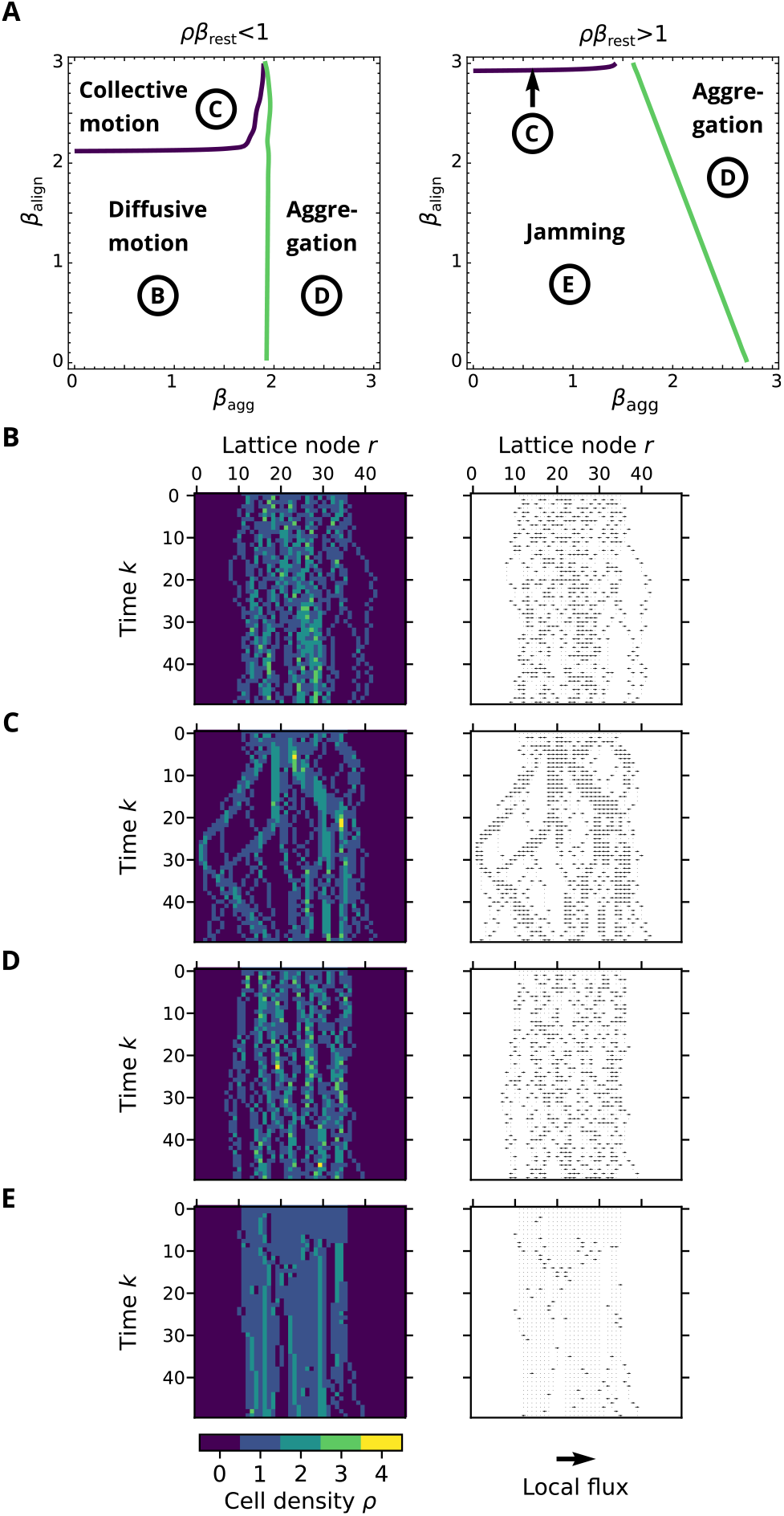
Migration modes in the BIO-LGCA model. (A) Phase diagram for low (left) and high (right) values of *β*_rest_. (B-E) Snapshots of the different phases of cell density (left) and local flux (right): (B) diffusive motion, if all sensitivities *β* are low: they interact weakly with one another and with the ECM; (C) collective motion for high *β*_align_; (D) pattern formation for high *β*_agg_; (E) jammed state for high *β*_rest_.

1. Diffusive (gas-like) phase. In this region, cells move freely and diffuse within the ECM duct. In the phase space, this corresponds to the region of the parameter space with low *β*_align_, *β*_agg_, and *β*_rest_ values, see Fig. 7B.
2. Collective motionn (active nematic) phase. This is the region with high *β*_align_. In this region, cells move collectively into the same direction, see Fig. 7C.
3. Aggregation phase. In this region, cells arrange themselves in static clusters and form cellular patterns within the duct. This corresponds to the region with high *β*_agg_., see Fig. 7D.
4. Jammed (glass-like) phase. This corresponds to the regime with high *β*_rest_. In this region, cells neither form patterns nor do they move collectively. However, almost all cells are in a resting channel and the dynamics is frozen, similar to a crystalline solid, see Fig. 7E.

In conclusion, we can reproduce the prime modes of collective migration using three mechanisms: alignment of velocities, aggregation and an inhibition of motion by non-migratory cells. We here treated these mechanisms as independent, however, in reality, they are the result of a complex interplay of cell-cell adhesion, cellular contractility, and intercellular signaling. Identifying the regulation of these cellular properties is a topic of ongoing research.

## 6 Discussion

In contrast to “continuum systems” and their canonical description with partial differential equations, there is no standard model for describing interactions of discrete objects, particularly interacting and migrating discrete biological cells. In this paper, the BIO-LGCA is proposed as a lattice-based model class for a spatially extended system of interacting cells. We provided various examples of BIO-LGCA models for homogeneous cell populations, i.e. cells are assumed to be of the same phenotype and not to change their behaviour. Examples include haptotaxis, chemotaxis, contact guidance and collective migration. However, the BIO-LGCA idea can be expanded to heterogeneous populations and environments, e.g. cells may dynamically regulate their adhesivities and/or may interact with a heterogeneous non-cellular environment [33]. BIO-LGCA models have already been used in the study of several biological processes including angiogenesis [34], bacterial rippling [35], active media [36], epidemiology [37] and various aspects of tumour dynamics [33, 29, 38, 39, 40, 16].

BIO-LGCA models are appropriate for low and moderate cell densities. For higher densities, e.g. in epithelial tissues, cell shape may matter and other models, such as the Cellular Potts model, might be better choices (see [2, 41] for reviews of on- and off-lattice models). It is also important to be aware of lattice artefacts inherent to the spatial discretisation inherent in every cellular automaton model, e.g. the checkerboard artefact (cp. [42]). In two spatial dimensions, the hexagonal lattice possesses less artefacts than the square lattice. A major advantage of BIO-LGCA models compared to other on- and off-lattice cell-based models for interacting cell systems, such as interacting particle systems, e.g. [43], asynchronous cellular automata, e.g. [44, 45, 46], further cell-based models [47] or systems of stochastic differential equations [48], is their computational efficiency, and their synchronicity and explicit velocity consideration, which enables the modelling of moderately packed cell collectives while minimizing model artefacts.

The BIO-LGCA framework facilitates theoretical analysis of collective behaviour [42]. In many cases, the collective behaviour of the BIO-LGCA can be analysed using reasonable approximations, such as a spatial mean-field description based on a partial differential equation [42, 49, 50, 51] (see also sec. 7). Collective behaviour at the tissue scale includes cell density patterns and the dynamics of moving cell fronts [52, 53]. Cell density patterns can often be assessed experimentally and provide, therefore, a means to relate BIO-LGCA model predictions to experimental observations.

Meanwhile, off-lattice models formulated as stochastic differential equations for moving and interacting cells have been derived from individual-based BIO-LGCA models [54]. Individual-based BIO-LGCA are lattice-based models that allow to distinguish and track individual cells which is not possible in classical LGCA models (e.g. [55]). We have also defined a “boson-type” LGCA which contrary to the BIO-LGCA does not possess an exclusion principle with respect to the velocity channels but facilitates mathematical analysis [56]. To model cell migration involving steric effects, the model has to be extended to discourage individuals moving into the direction of steeply increasing cell density gradients.

The BIO-LGCA modelling strategy is “modular”: starting from “basic model moduls”, which include those explored in this paper such as alignment, attraction, contact guidance, hapto- and chemotaxis. Coupling them is required to design models for complex biological problems. The focus of future activities is the analysis of further model combinations for selected biological problems, which are not necessarily restricted to cells but could also comprise interactions at subcellular and tissue scales. The resulting multi-scale models will contain a multitude of coupled spatial and temporal scales and will impose significant challenges for their analytic treatment.

## 7 Acknowledgments

HH gratefully acknowledges the funding support of the Helmholtz Association of German Research Centers—Initiative and Networking Fund for the project on Reduced Complexity Models (ZT-I-0010). HH is supported by MulticellML (01ZX1707C) of the Federal Ministry of Education and Research (BMBF) and the Volkswagenstiftung within the “Life?” programm (96732). AD acknowledges support by the EU-ERACOSYS project no. 031L0139B. JMNS acknowledges support from the PAPIIT-UNAM grant, project IA104821. SS is supported by the European Social Fund (ESF), co-financed by tax funds based on the budget adopted by the members of the Saxon State Parliament. The authors thank the Centre for Information Services and High Performance Computing at TU Dresden for providing an excellent infrastructure.

## Footnotes

1 𝒩 ^*b*^ + **r** = {**r** + **r** *′*, **r** *′* ∈ *𝒩* ^*b*^}

## References

[1] F. Bertaux, S. Stoma, D. Drasdo, and G. Batt, “Modeling dynamics of cell-to-cell variability in trail-induced apoptosis explains fractional killing and predicts reversible resistance,” PLoS Comput Biol, vol. 10, no. 10, p. e1003893, 2014.

[2] P. V. Liedekerke, M. M. Palm, N. Jagiella, and D. Drasdo, “Simulating tissue mechanics with agent-based models: concepts, perspectives and some novel results,” Computat. Part. Mechan., vol. 2, pp. 401–444, 2015.

[3] M. J. Plank and M. J. Simpson, “Models of collective cell behaviour with crowding effects: comparing lattice-based and lattice-free approaches,” Journ. Roy. Soc. Interface, vol. 9, p. 2983–2996, 2012.

[4] S. T. Johnston, M. J. Simpson, and M. J. Plank, “Lattice-free descriptions of collective motion with crowding and adhesion,” Phys. Rev. E, vol. 88, p. 062720, 2013.

[5] T. J. Newman and R. Grima, “Many-body theory of chemotactic cell-cell interactions,” Physic. Rev. E, vol. 70, p. 051916, 2004.

[6] A. M. Middleton, C. Fleck, and R. Grima, “A continuum approximation to an off-lattice individual-cell based model of cell migration and adhesion,” J. Theoret. Biol., vol. 359, pp. 220–232, 2014.

[7] O. M. Matsiaka, C. J. Penington, R. E. Baker, and M. J. Simpson, “Continuum approximations for lattice-free multispecies models of collective cell migration,” Journal of Theoretical Biology, vol. 422, pp. 1–11, 2017.

[8] R. Grima and S. Schnell, “A mesoscopic simulation approach for modeling intracellular reactions,” J. Statistic. Phys., vol. 128, pp. 139–164, 2007.

[9] M. Radszuweit, M. Block, J. Hengstler, E. Schöll, and D. Drasdo, “Comparing the growth kinetics of cell populations in two and three dimensions,” Physical Review E, vol. 79, no. 5, p. 051907, 2009.

[10] F. Graner and J. A. Glazier, “Simulation of biological cell sorting using a two-dimensional extended potts model,” Physical review letters, vol. 69, no. 13, p. 2013, 1992.

[11] A. W. Burks, Essays on Cellular Automata. University of Illinois Press, Urbana IL, 1970.

[12] J. L. Casti, Alternate realities. New York: John Wiley, 1989.

[13] B. Chopard and M. Droz, Cellular automata modeling of physical systems. Cambridge University Press, New York, 1998.

[14] S. Wolfram, A new kind of science. Wolfram Media, Inc, 2002.

[15] E. Gavagnin and C. A. Yates, “Modeling persistence of motion in a crowded environment: The diffusive limit of excluding velocity-jump processes,” Physical Review E, vol. 97, no. 3, p. 032416, 2018.

[16] O. Ilina, P. Gritsenko, S. Syga, A. Deutsch, and P. Friedl, “Cell–cell adhesion and 3d matrix confinement determine jamming transitions in breast cancer invasion,” Nat Cell Biol, vol. 22, p. 1103–1115, 2020.

[17] J. M. Nava-Sedeño, H. Hatzikirou, F. Peruani, and A. Deutsch, “Extracting cellular automaton rules from physical Langevin equation models for single and collective cell migration,” Journal of mathematical biology, vol. 75, no. 5, pp. 1075–1100, 2017.

[18] R. Grima and S. Schnell, “A mesoscopic simulation approach for modeling intracellular reactions,” Journal of Statistical Physics, vol. 128, no. 1, pp. 139–164, 2007.

[19] J. M. Nava-Sedeño, H. Hatzikirou, R. Klages, and A. Deutsch, “Cellular automaton models for time-correlated random walks: derivation and analysis,” Scientific reports, vol. 7, no. 1, pp. 1–13, 2017.

[20] K. Kawasaki, “Simple derivations of generalized linear and nonlinear Langevin equations,” Journal of Physics A: Mathematical, Nuclear and General, vol. 6, pp. 1289–1295, sep 1973.

[21] F. Peruani, M. Bär, and A. Deutsch, “A mean-field theory for self-propelled particles interacting by velocity alignment mechanisms,” Eur. Phys. J., vol. 157, no. 111, 2008.

[22] P. Romanczuk, M. Bär, W. Ebeling, B. Lindner, and L. Schimansky-Geier, “Active brownian particles: From individual to collective stochastic dynamics,” Europ. Phys. J. Spec. Top., vol. 202, no. 1, pp. 1–162, 2012.

[23] H. Takagi, M. J. Sato, T. Yanagida, and M. Ueda, “Functional analysis of spontaneous cell movement under different physiological conditions,” PloS one, vol. 3, no. 7, p. e2648, 2008.

[24] S. Pressé, K. Ghosh, J. Lee, and K. A. Dill, “Principles of maximum entropy and maximum caliber in statistical physics,” Reviews of Modern Physics, vol. 85, no. 3, p. 1115, 2013.

[25] R. B. Dickinson and R. T. Tranquillo, “A stochastic model for cell random motility and haptotaxis based on adhesion receptor fuctuations,” J. Math. Biol., vol. 31, pp. 563–600, 1993.

[26] H. Bussemaker, A. Deutsch, and E. Geigant, “Mean-field analysis of a dynamical phase transition in a cellular automaton model for collective motion,” Phys. Rev. Lett., vol. 78, pp. 5018–5021, 1997.

[27] H. J. Bussemaker, “Analysis of a pattern forming lattice-gas automaton: mean-field theory and beyond,” Phys. Rev. E, vol. 53, no. 2, pp. 1644–1661, 1996.

[28] A. R. Kansal, S. Torquato, G. R. Harsh, E. A. Chiocca, and T. S. Deisboeck, “Simulated brain tumor growth using a three-dimensional cellular automaton,” J. Theor. Biol., vol. 203, p. 367, 2000.

[29] J. Moreira and A. Deutsch, “Cellular automaton models of tumour development – a critical review,” Adv. Compl. Syst. (ACS), vol. 5, no. 2, pp. 1–21, 2002.

[30] P. Friedl, J. Locker, E. Sahai, and J. E. Segall, “Classifying collective cancer cell invasion,” Nature cell biology, vol. 14, no. 8, pp. 777–783, 2012.

[31] M. Sadati, N. T. Qazvini, R. Krishnan, C. Y. Park, and J. J. Fredberg, “Collective migration and cell jamming,” Differentiation, vol. 86, no. 3, pp. 121–125, 2013.

[32] A. G. Clark and D. M. Vignjevic, “Modes of cancer cell invasion and the role of the microenvironment,” Current opinion in cell biology, vol. 36, pp. 13–22, 2015.

[33] D. Reher, B. Klink, A. Voss-Boehme, and A. Deutsch, “Cell adhesion heterogeneity reinforces tumour cell dissemination: novel insights from a mathematical model,” Biol. Direct, 2017.

[34] C. Mente, I. Prade, L. Brusch, G. Breier, and A. Deutsch, “Parameter estimation with a novel gradient-based optimization method for biological lattice-gas cellular automaton models,” J. Math. Biol., vol. 63, no. 1, pp. 173–200, 2010.

[35] M. Alber, M. Kiskowski, and Y. Jiang, “Lattice gas cellular automata model for rippling in myxobacteria,” Physica D, vol. 191, p. 343, 2004.

[36] S. Syga, J. M. Nava-Sedeño, L. Brusch, and A. Deutsch, “A lattice-gas cellular automaton model for discrete excitable media,” in Spirals and Vortices, pp. 253–264, Springer, 2019.

[37] H. Fuks and A. T. Lawniczak, “Individual-based lattice model for spatial spread of epidemics,” Discrete Dynamics in Nature and Society, vol. 6, 2001.

[38] K. Böttger, H. Hatzikirou, A. Voß-Böhme, E. A. Cavalcanti-Adam, M. A. Herrero, and A. Deutsch, “An emerging Allee effect is critical for tumor initiation and persistence,” PLoS Comput. Biol., vol. 11, no. 9, p. e1004366, 2015.

[39] H. Hatzikirou, L. Brusch, C. Schaller, M. Simon, and A. Deutsch, “Prediction of traveling front behavior in a lattice-gas cellular automaton model for tumor invasion,” Comput. Mathem. Applic., vol. 59, pp. 2326–2339, April 2010.

[40] M. Tektonidis, H. Hatzikirou, A. Chauviere, M. Simon, K. Schaller, and A. Deutsch, “Identification of intrinsic in vitro cellular mechanisms for glioma invasion,” J. Theor. Biol., vol. 287, pp. 131–147, October 2011.

[41] T. Boekhorst, L. Preziosi, and P. Friedl, “Plasticity of cell migration in vivo and in silico,” Annu. Rev. Cell Dev. Biol., vol. 32, pp. 491–526, 2016.

[42] A. Deutsch and S. Dormann, Cellular automaton modeling of biological pattern formation. Birkhauser. 334 p, 2005, 2018 (2nd ed.).

[43] T. M. Liggett, Interacting Particle Systems. New York: Springer, 1985.

[44] M. Badoual, C. Deroulers, M. Aubert, and B. Grammaticos, “Modelling intercellular communication and its effects on tumour invasion,” Physical biology, vol. 7, no. 4, p. 046013, 2010.

[45] B. J. Binder, K. A. Landman, D. F. Newgreen, J. E. Simkin, Y. Takahashi, and D. Zhang, “Spatial analysis of multi-species exclusion processes: application to neural crest cell migration in the embryonic gut,” Bulletin of mathematical biology, vol. 74, no. 2, pp. 474–490, 2012.

[46] J. Bloomfield, J. Sherratt, K. Painter, and G. Landini, “Cellular automata and integrodifferential equation models for cell renewal in mosaic tissues,” Journal of the Royal Society Interface, vol. 7, no. 52, pp. 1525–1535, 2010.

[47] J. Galle, M. Hoffmann, and G. Aust, “From single cells to tissue architecture—a bottom-up approach to modelling the spatio-temporal organisation of complex multi-cellular systems,” Journal of mathematical biology, vol. 58, no. 1-2, p. 261, 2009.

[48] K. A. Rejniak and A. R. Anderson, “Hybrid models of tumor growth,” Wiley Interdisciplinary Reviews: Systems Biology and Medicine, vol. 3, no. 1, pp. 115–125, 2011.

[49] U. Frisch, B. Hasslacher, and Y. Pomeau, “Lattice-gas automata for the navier-stokes equation,” Phys. Rev. Lett., vol. 56, no. 14, pp. 1505–1508, 1986.

[50] D. Wolf-Gladrow, Lattice-gas cellular automata and lattice Boltzmann models: an introduction. New York: Springer, 2000.

[51] S. Wolfram, Cellular Automata and Complexity - collected papers. Addison-Wesley, 1994.

[52] H. Hatzikirou, L. Brusch, and A. Deutsch, “From cellular automaton rules to an effective macroscopic mean-field description,” Acta Phys. Polon. B Proc. Suppl., vol. 3, pp. 399–416, 2010.

[53] C. Mente, I. Prade, L. Brusch, G. Breier, and A. Deutsch, “A lattice-gas cellular automaton model for in vitro sprouting angiogenesis,” Acta Phys. Pol. B, vol. 5, no. 1, pp. 99–115, 2012.

[54] C. Mente, A. Voß-Böhme, and A. Deutsch, “Analysis of individual cell trajectories in lattice-gas cellular automaton models for migrating cell populations,” Bull. Mathem. Biol., vol. 77, no. 4, pp. 1–38, 2015.

[55] U. Frisch, B. Hasslacher, and Y. Pomeau, “Lattice-gas automata for the Navier-Stokes equation,” Phys. Rev. Lett., vol. 56, No. 14, pp. 1505–1509, 1986.

[56] J. M. Nava-Sedeno, H. Hatzikirou, A. Voß-Böhme, L. Brusch, A. Deutsch, and F. Peruani, “Vectorial active matter on the lattice: emergence of polar condensates and nematic bands in an active zero-range process.” archives ouvertes: hal-02460291, 2020.

